# Impaired lysine biosynthesis primes constitutive energy stress and dark-stress sensitivity

**DOI:** 10.1101/2025.09.10.675385

**Authors:** Débora Gonçalves Gouveia, Wesley E. B. Barrios, Philipp Westhoff, Samantha Flachbart, João Henrique F. Cavalcanti, Bruno P. Zanotti, Alisdair R. Fernie, Adriano Nunes-Nesi, Andreas P. M. Weber, Wagner L. Araújo

## Abstract

Plant survival under prolonged darkness relies on dynamic metabolic reprogramming that redirects limited carbon resources toward mitochondrial respiration and nutrient remobilization. Although amino acids function as alternative respiratory substrates during carbon starvation, how their biosynthesis integrates into this metabolic adjustment remains poorly understood. Here, we investigated the role of lysine biosynthesis using the *dapat* mutant, which exhibits photoperiod-dependent hypersensitivity to extended darkness. Under short-day conditions, *dapat* plants exhibited accelerated senescence. reduced photosystem II maximum quantum efficiency, pronounced protein degradation, and accumulation of amino acid, ultimately failing to recover after nine days of darkness. By contrast, survival was largely restored under neutral-day conditions, indicating that restricted carbohydrate reserves during short-day growth exacerbate the mutant phenotype. Remarkably, transcript profiling further revealed that reduced DAPAT activity constitutively activates a catabolic program, including genes associated with amino acid degradation, alternative respiratory pathways, senescence, and autophagy, even under non-stress conditions. Together, these findings identify DAPAT-mediated lysine biosynthesis as a central metabolic hub linking carbon availability to energy and stress signaling. Disruption of this pathway compromises metabolic flexibility and limits the capacity to withstand prolonged carbon starvation. Our results provide new insights into how primary metabolism coordinates stress responses to extended darkness and highlight amino acid biosynthesis as an important component of plant resilience under energy-deprived darkness conditions.

## 1. Introduction

Plant growth and survival depend on light as the primary source of energy. Extended periods of darkness impose severe physiological stress and trigger complex metabolic reprogramming that is essential for survival under carbon-limited heterotrophic conditions (Law *et al*., 2018). Extended darkness leads to carbohydrate starvation, prompting plants to shift from canonical carbon reserves to alternative energy sources, thereby fundamentally altering their metabolic priorities. In this rewired metabolic state, plant cells use a variety of cellular substrates to sustain ATP production (Baena-González *et al*., 2007; Brandt *et al*., 2018; Zhu *et al*., 2022). To feed the tricarboxylic acid (TCA) cycle during extended darkness, plants mobilize reduced carbon released from cell wall degradation, fatty acids released through β-oxidation (Kunz *et al*., 2009; Fan et al., 2017), and amino acids resulting from protein degradation (Hildebrandt, 2018). In parallel, electrons derived from amino acid catabolism are transferred to the mitochondrial electron transport chain (mETC) via the ubiquinol pool through the ETF/ETFQO system (Ishizaki *et al*., 2005; Araújo *et al*., 2010, 2011, 2012).

Lysine metabolism is a key regulatory node in plant adaptive responses (Singh et al., 2025), that mediates the crosstalk between the TCA cycle and mETC, which underlies respiratory processes and stress resilience (Araújo *et al*., 2010; Galili, 2011; Kirma *et al*., 2012; Yang *et al*, 2020). The importance of lysine is highlighted by the embryo-lethality of the *Arabidopsis* knockout mutant lacking the *L,L*-diaminopimelate aminotransferase (*DAPAT)* gene (AT4G33680), which encodes a crucial enzyme in lysine biosynthesis (Song *et al*., 2004; Hudson *et al*., 2005, 2006). Even under optimal conditions, disruptions in lysine synthesis induce chronic energy deprivation, characterized by impaired growth, massive metabolic reprogramming and altered stress sensitivity (Cavalcanti *et al*., 2018; Neves *et al*., 2024; Gouveia *et al*., 2026). Together, these features point to an extensive uncoupling of carbon and nitrogen metabolism, likely intensified by compensatory mitochondrial pathways (Araújo *et al*., 2010, 2011; Cavalcanti *et al*., 2018).

In this context, we propose that lysine biosynthesis functions as a photoperiod-sensitive metabolic hub that coordinates carbon status, amino acid catabolism, and mitochondrial respiration during prolonged darkness. Using the *Arabidopsis* lysine-biosynthesis mutant *dapat* under contrasting photoperiods, we show that reduced DAPAT activity establishes a pre-stressed metabolic state that becomes particularly severe when carbohydrate reserves are limited, leading to accelerated senescence and impaired recovery after darkness. In short-day plants, this defect is exacerbated, whereas neutral-day growth partially mitigates metabolic failure and allows survival after stress release. These results position lysine homeostasis as an important regulator of metabolic flexibility during carbon starvation and extend previous observations on the *dapat* mutant.

## 2. Materials and methods

### 2.1. Plant material

*Arabidopsis* ecotype Col-0 the *dapat* mutant - a lysine biosynthesis-deficient line (Rate & Greenberg, 2001; Song *et al*., 2004; Hudson *et al*., 2006; Cavalcanti *et al*., 2018; Neves *et al*., 2024; Gouveia *et al*., 2026), were used in this study. *Arabidopsis* seeds were surface-sterilized by stirring in 1 mL of 80% (v/v) ethanol containing 0.05% (v/v) Tween 20 for 5 minutes, followed by a 5-minute wash in 100% ethanol. Residual ethanol was removed, and seeds were air-dried for 2 hours under sterile conditions. Sterilized seeds were grown on half-strength Murashige and Skoog (Murashige & Skoog, 1962) medium (pH 5.7) with sucrose supplemented 1% (w/v) and 0.8% (w/v) agar. Plates were sealed with micropore tape and stratified in darkness at 4°C for 72 hours to synchronize germination.

### 2.2. Growth conditions and dark treatment

#### 2.2.1. Short-day conditions

Seeds were germinated at 20°C under short-day conditions (8h/16h light-dark), 60% relative humidity, and 150 µmol photons m^−2^s^−1^ light intensity. For the dark treatment, 12-day-old seedlings were transferred to pots and grown at 20°C (day) and 18°C (night), under the previously described conditions. Four-week-old plants were transferred to darkness in the same growth cabinet, where they were kept for 9 days. Whole rosettes were harvested at 0, 3, 5, 7, and 9 days after transition to darkness, immediately frozen in liquid nitrogen, and then stored at 80°C until further analysis. Five biological replicates were used per genotype: treatment group.

#### 2.2.2. Neutral-day conditions

Seeds were germinated at 20°C under neutral-day conditions (12h/12h light–dark), 60% relative humidity, and 150 µmol photons m ² s ¹ light intensity. For the dark treatment, 12-day-old seedlings were transferred to soil and grown at 20°C (day) and 18°C (night), under the previously described conditions. Four-week-old plants were transferred to darkness in the same growth cabinet, where the treatment was imposed for 9 days, followed by a 4-day recovery period under initial growth conditions. Whole rosettes were collected at 0, 3, 5, 7, and 9 days after the transition to darkness, as well as during the recovery phase. Five biological replicates were used per genotype: treatment group.

### 2.3. Measurements of respiration and photosynthetic parameters

Dark respiration measurements were performed on fully expanded leaves of 5-week-old plants maintained in darkness using an open-flow infrared gas-exchange analyzer system (LICOR 6400XT). The *Fv/Fm* ratio, an indicator of the maximum potential quantum efficiency of PSII, was measured according to established protocol (Oh *et al*., 1996).

### 2.4. Determination of metabolite levels

Methanolic extraction was performed using approximately 25 mg of liquid nitrogen-ground fresh leaf tissue, followed by immediate addition of extraction buffer. Chlorophyll content (*a* and *b*) were determined as previously described (Porra *et al*., 1989). Total protein level was measured using the insoluble fraction according to (Bradford, 1976). In the soluble fraction, total soluble amino acids were quantified following (Gibon *et al.,* 2004).

### 2.5. Metabolite profiling

Metabolites were extracted using 1.5 mL of pre-cooled (−20°C) mixture of H_2_O:MeOH:CHCl_3_ (1:2.5:1 v:v) extraction buffer was used, supplemented with 50 μL/5 mM Ribitol/DMPA internal standards solution per 50 mL buffer. Sample were mixed by vortexed for 20 sec. Samples were shaken on an orbital shaker for 10 min at 4°C. The samples were then centrifuged at 20,000 *g* for 5 min at 4°C. After centrifugation, 1.0 mL of supernatant was carefully aspirated and transferred to a clean 1.5-ml microcentrifuge tube. Samples were stored at -80°C until derivatization. For GC-MS analysis 30 μl of sample was freeze dried using speed-vac and derivatized (Gu *et al*., 2012; Shim *et al*., 2019). Raw data were converted to the mzXML format using ProteoWizard and to the NetCDF format via MetAlign using default parameters. Deconvolution of mass spectra was conducted using the free deconvolution software AMDIS (Automated Mass Spectral Deconvolution and Identification System from NIST), and metabolites were identified using the NIST14 Mass Spectral Library (https://www.nist.gov/srd/niststandard-reference-database-1a-v14). Database matches with more than 70% were compared and validated with an in-house chemical standard library for compound annotation. Unidentified compounds are assigned a name that combines their presumed chemical family and their specific retention time. Extracted peaks were integrated using MassHunter Quantitative (v b08.00, Agilent Technologies), all metabolite levels were normalized to fresh weight and Ribitol internal standard (Sigma-Aldrich) to correct technical errors.

### 2.6. Gene expression analysis

Total RNA was extracted with TRIzol reagent (Ambion, Life Technology) from five individual plants per genotype/treatment. After purification, RNA concentration was quantified using a NanoDrop ND-2000 spectrophotometer (Thermo Scientific NanoDrop Technologies, Wilmington, Delaware, United States) and integrity verified on a 1,5 % (w/v) agarose gels. A total of 0.5 μg of RNA was treated with DNAse I (Invitrogen®) to remove potential contamination with genomic DNA. cDNA synthesis was performed using 0.5 μg of total RNA from the four highest-quality samples per genotype/treatment with GoScript™ Reverse Transcriptase (Afive003), RNasin Ribonuclease Inhibitor, and Oligo(dT) (A2791) following manufacturer protocol. For gene expression analysis, Ludwig qPCR SYBR-green mix/ROX (ID:40) was used following the manufacturer protocol with MicroAmpTM Optical 96-well Reaction Plate (Applied Biosystems, Singapore, China) and MicroAmpTM Optical Adhesive Film (Applied Biosystems, Foster City, CA, USA). Primers used for RT-qPCR (Table S1) were designed using Primer3Plus (Untergasser *et al*., 2012). Gene expression analyses were performed using four replicates per genotype/treatment with a minimum of three replicates analyzed due to occasional technical issues, and relative expression was calculated by the 2^-(ΔΔCt)^ method, presented as mean ± standard error for each group. Expression levels were normalized using the reference genes: *UBC9* and *UBC21* (short-day experiment) and *MON1* and *TIP41* (neutral-day experiment).

### 2.7. Experimental design and statistical analysis

All statistical analyses were performed in R version 4.5.0 using the stats, ggplot2, and heatmap packages. Two-tailed Student’s *t*-tests were applied with the significance threshold set at *p* < 0.05; values with 0.05 < p < 0.1 were indicated in boxplots. Significant differences in heatmaps were denoted by asterisks at *p* < 0.05. Comparisons were made using wild-type day 0 as the reference group for the ΔΔCt method but compared statistically against WT at each day, except for *DAPAT* gene expression, which used wild-type: day 0 as the reference group for both gene expression assessment and comparison. Heatmaps present wild-type Control: day 0 normalized log -fold changes for all the metabolites dataset, with two-tailed Student’s *t*-tests used to each metabolite (row effects) and significant differences set at *p* < 0.05 evidenced directly on heatmap cells by an asterisk.

## 3. Results

### Darkness reveals the metabolic cost of lysine deficiency

Building on the previously reported senescence phenotype of the *Arabidopsis* lysine biosynthesis-deficient *dapat* mutant (Cavalcanti *et al*., 2018), confirmed by independent complementation lines grown under standard conditions (Figure S1), we hypothesized that lysine deficiency compromises the metabolic coordination required to integrate amino acid catabolism, respiratory metabolism and growth control during prolonged darkness. To test this hypothesis, we subjected wild-type (WT) and *dapat* plants to nine days of continuous darkness. Throughout the treatment, mutant plants displayed accelerated senescence (Figure S2A, day 5) and failed to recover after re-exposure to light (Figure S2B), indicating a marked sensitivity of *dapat* to prolonged carbon starvation.

To further examine the metabolic and molecular consequences of extended darkness, an independent set of WT and *dapat* plants was also subjected to the same dark treatment for sampling (Figure 1A, B). We also monitored two key indicators of chloroplast function, the maximum quantum efficiency of photosystem II (*Fv/Fm*) (Figure 1C) and total chlorophyll content (Figure S3). In WT plants, *Fv/Fm* remained relatively stable throughout the 9-day treatment, with only minor temporal declines (Figure 1C). By contrast, *dapat* displayed a constitutively lower *Fv/Fm* even under control conditions, consistent with a pre-existing impairment in *dapat* photosynthetic performance linked to lysine biosynthesis deficiency, and this difference became more pronounced under darkness. By day 7, PSII efficiency had declined significantly more in *dapat* than in WT plants (Figure 1C). Along with this loss of photochemical capacity, total chlorophyll content and the chlorophyll a/b ratio decreased progressively during darkness, with stronger reductions in *dapat* than in WT (Figure S3C). Notably, the lower chlorophyll content and reduced *Fv/Fm* were already evident in the mutant plants before stress imposition, indicating that *dapat* plants enter darkness with an intrinsic photosynthetic disadvantage.

**Figure 1.**
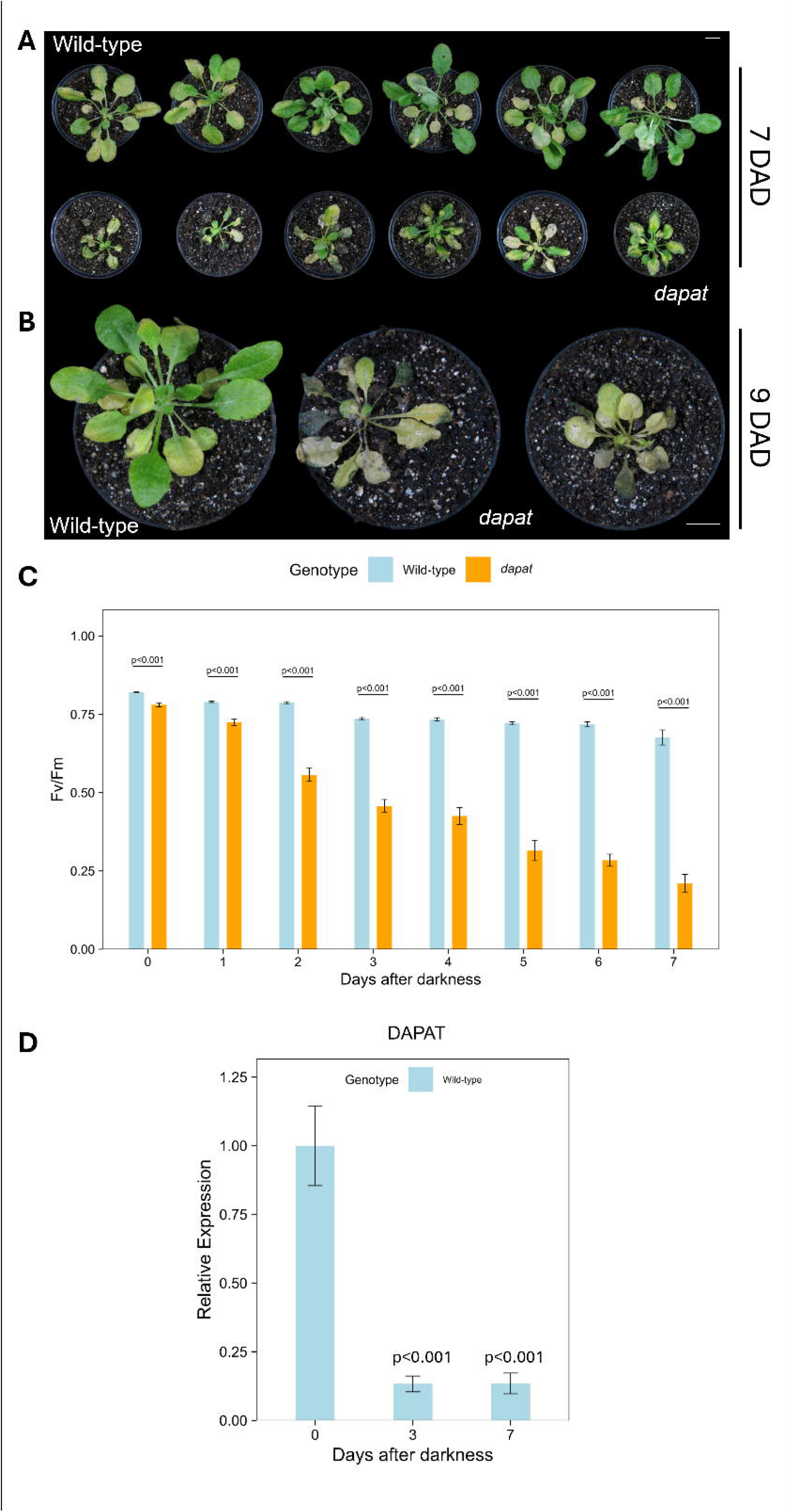
The lysine biosynthesis-deficient *dapat* mutant is hypersensitive to extended darkness and displays impaired metabolic recovery. (A) Representative phenotypes of five-week-old *Arabidopsis thaliana* wild-type (WT) and *dapat* mutant plants on day 7, of the 9-day dark treatment (DAD). Plants were initially grown under short-day (8h/16h light-dark) conditions, 150 μmol photons m ²s ¹, 20°C day/18°C night for five weeks before exposure to continuous darkness. (B) Comparative phenotypic analysis of five-week-old plants after nine days after darkness (DAD). (C) Maximum quantum efficiency of photosystem II (*Fv/Fm*) in leaves of five-week-old plants during 7-days of extended darkness (n = 8). (D) RT-qPCR analysis of *DAPAT* (AT4G33680) transcript levels, under control conditions (day 0) and after exposure to extended darkness (days 3, and 7). Expression levels are presented as log fold change relative to day 0 and were measured at the end of the light period. Data represent means ± SEM of three biological replicates. P values indicate significant differences determined by two-tailed Student’s t-test between WT and *dapat* at each time point. Expression levels were normalized using the reference genes *UBC9* and *UBC21*. The reference group for the ΔΔCt method used was WT: day 0. White scale bars at the right top and bottom represent 1 cm.

Given the intrinsic disruption of carbon–nitrogen homeostasis and increased amino acid pools in *dapat* plants (Cavalcanti *et al*., 2018; Gouveia *et al*., 2026), we next investigated changes in protein and amino acid pools during prolonged darkness. This analysis revealed genotype-specific responses (Figure 2). Total protein decreased in both genotypes after day 5 of darkness (Figure 2A), whereas amino acid pools increased throughout the treatment (Figure 2B), consistent with enhanced proteolysis under starvation. Notably, after 9 days of darkness, *dapat* plants accumulated significantly more amino acids than WT (Figure 2B), indicating a defective nitrogen remobilization or sustained C/N imbalance under prolonged stress. Together, these data suggest that dark-induced senescence in *dapat* reflects more than simple energy deprivation; instead, it likely arises from a broader failure to coordinate lysine homeostasis with the metabolic and developmental processes that sustain growth during starvation, consistent with the view that lysine biosynthesis acts as a metabolic checkpoint in *Arabidopsis* (Gouveia *et al*., 2026).

**Figure 2.**
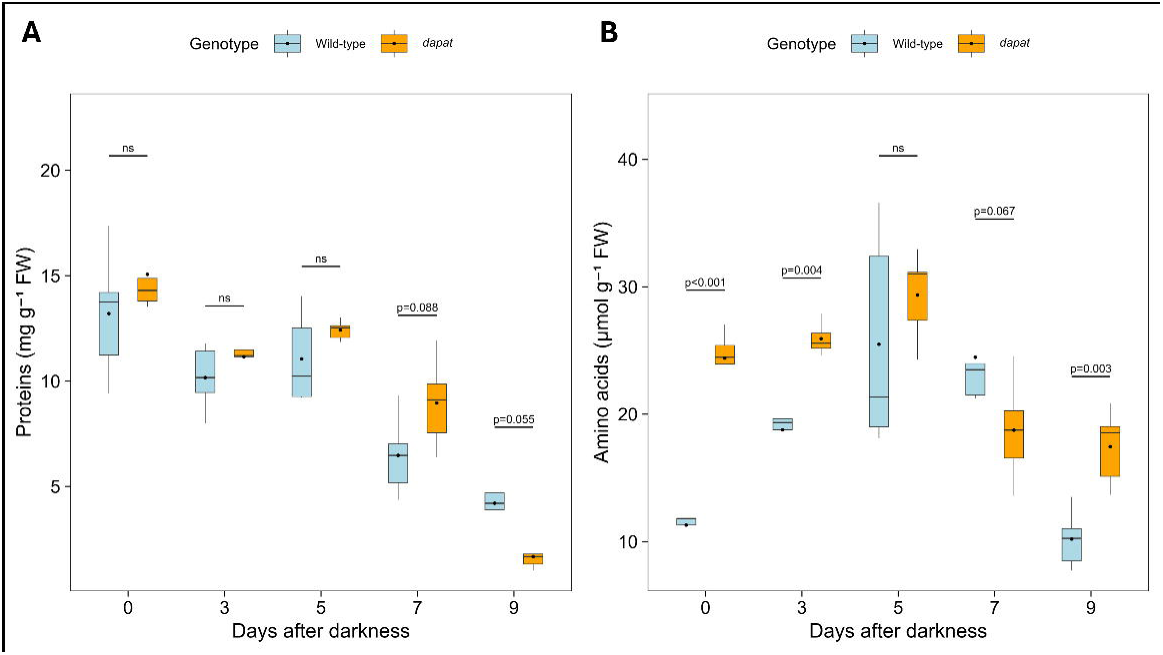
Lysine biosynthesis deficiency intensifies amino acid accumulation during dark-induced senescence. (A) Total proteins and (B) amino acids content in leaves of five-week-old *Arabidopsis thaliana* wild-type (WT) and *dapat* mutant plants grown under short-days (8h/16h light-dark) conditions after 0, 3, 5, 7 and 9 days under extended darkness. P values indicate significant differences determined by two-tailed Student’s t-test between WT and *dapat* at each time point. Values represent the mean ± standard error of five independent samples.

### DAPAT activity sets the metabolic threshold for survival in darkness

The premature senescence phenotype of *dapat* plants, together with their reduced survival under prolonged darkness (Figure S2B), prompted us to examine whether *DAPAT* expression is regulated during extended carbon deprivation. In line with the well-established transcriptional response of plants to prolonged darkness, in which biosynthetic pathways are repressed while autophagy and other catabolic processes are activated to expand the pool of alternative respiratory substrates (Barros *et al*., 2017; Hirota *et al*., 2018), DAPAT transcript levels in WT plants were strongly downregulated during the treatment, showing a 7.4-fold decrease by day 7 relative to the control (day 0; Figure 1D). This coordinated metabolic reprogramming likely reflects the adaptive shift triggered by suppressed carbon assimilation during extended darkness (Baena-González *et al*., 2007; Singh *et al*., 2009; Liu & Bassham, 2012; Marshall *et al*., 2018; Hildebrandt, 2018).

On this basis, we next investigated amino acid catabolism and mitochondrial alternative respiratory pathways, given the critical role of amino acids as electron donors to the mitochondrial electron transport chain during energy deprivation (Araújo *et al*., 2010; Schertl & Braun, 2014; Hirota *et al*., 2018). Expression levels highlighted distinct regulation patterns for genes encoding branched-chain amino acid transferase 2 (BCAT-2) (AT1G10070), D-2-hydroxyglutarate dehydrogenase (D2HGDH) (AT4G36400), isovaleryl-CoA dehydrogenase (IVDH) (AT1G04710), electron transfer flavoprotein β subunit (ETFβ) (AT2G32360), and ETF: ubiquinone oxidoreductase (ETFQO) (AT2G43400) (Figure 3A–E). *BCAT-2* expression exhibited subtle upregulation in *dapat* plants at day 0, with both genotypes showing strong induction that peaked on day 3 (∼60-fold in WT and ∼45-fold in *dapat*) before declining by day 7 (Figure 3A), accompanied by a marked downregulation in mutant plants. Conversely, *IVDH* transcript levels displayed subtle downregulation in *dapat* plants under control conditions (Figure 3B), with significant and sustained induction in the mutant throughout the treatment period, suggesting that a potential increase of branched-chain amino acid (BCAAs) degradation pathway under extended darkness compensates for the impaired lysine biosynthesis.

**Figure 3.**
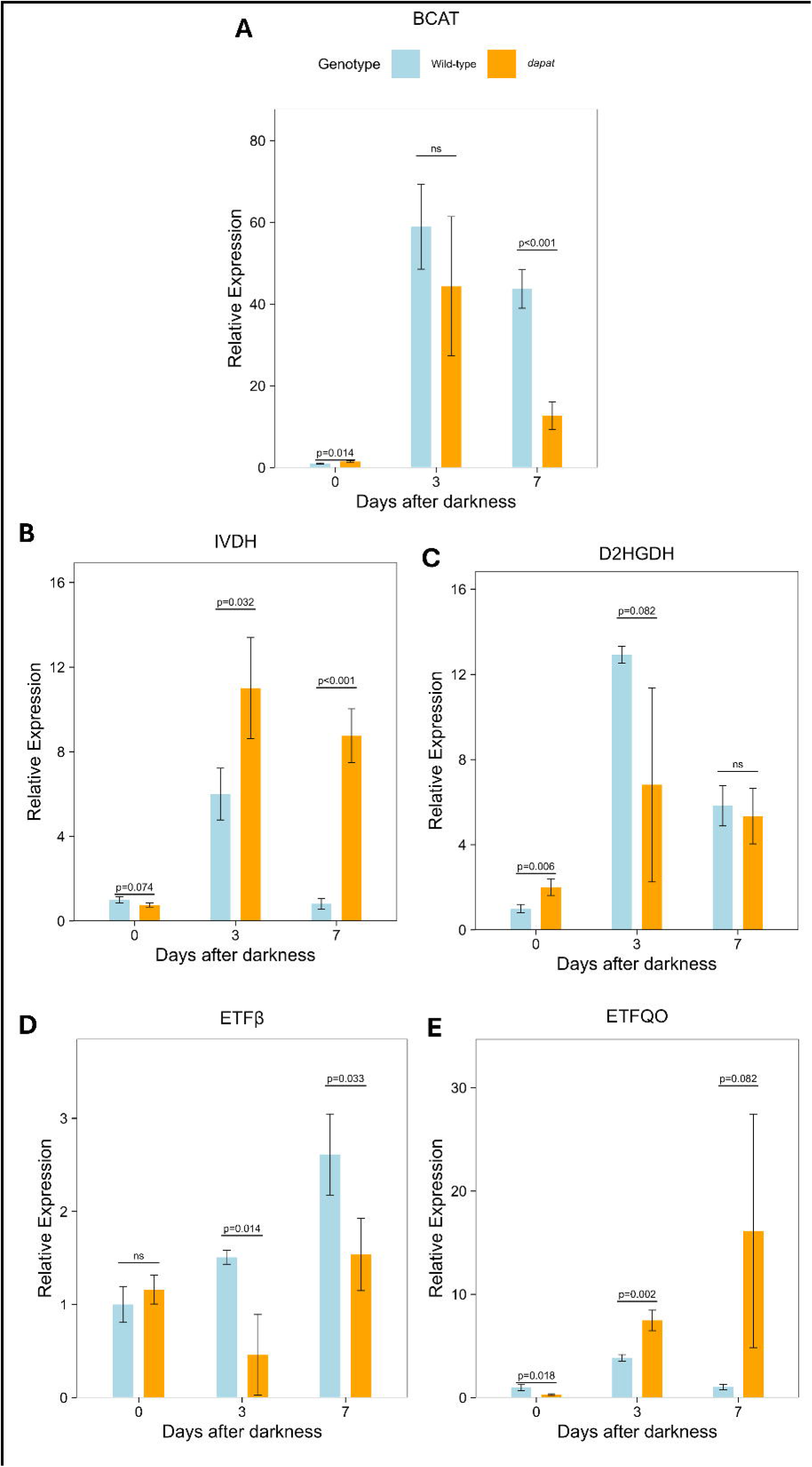
Extended darkness differentially reprograms alternative respiratory pathway gene expression in the *dapat* mutant. Transcript levels of (A) *BCAT-2*, (B) *IVDH*, (C) *D2HGDH*, (D) *ETF*β, and (E) *ETFQO* in *Arabidopsis thaliana* wild-type (WT) and *dapat* mutant plants grown under short-day (8h/16h light-dark) conditions, 150 μmol photons m ²s ¹, 20°C day/18°C night and subjected to extended darkness. Data represent transcript abundance normalized to wild-type at day 0 (dashed line) and are expressed as mean ± SEM of three biological replicates. P values indicate significant differences versus WT at each time point by two-tailed Student’s t-test. Expression levels were normalized using the reference genes *UBC9* and *UBC21*. The reference group for the ΔΔCt method used was WT: day 0.

Further examination of lysine catabolism and downstream metabolic pathways revealed distinct genotype-dependent regulation of *D*2HGDH. In *dapat*, *D*2HGDH showed a strong constitutive upregulation from day 0, whereas WT displayed a sharp transient induction peaking at day 3, with a 13-fold increase (Figure 3C); by day 7, transcript levels in both genotypes converged to similar values. Components of the ETF/ETFQO complex also showed divergent regulation. In WT, *ETF*β expression progressively increased, reaching 2.6-fold by day 7, whereas it remained suppressed in *dapat* (Figure 3D). By contrast, *ETFQO*, which was initially downregulated in *dapat* under control conditions (day 0), became strongly induced during dark-induced senescence, reaching a 16-fold increase by day 7, while remaining low in WT plants (Figure 3E). Together, these findings suggest that WT transiently mobilizes lysine-derived substrates as an adaptive response to darkness, whereas *dapat* plants display constitutively elevated expression of catabolic targets, likely reflecting an underlying metabolic dysfunction, followed by an attenuated response throughout the treatment period.

### Lysine deficiency rewires starvation responses beyond alternative respiration

Following the altered metabolic and senescence-related responses observed in dapat under extended darkness, we next assessed autophagy- and senescence-associated programs by analyzing the relative expression of autophagy-related genes *ATG7* (AT5G46830), *ATG8b* (AT4G04620), and *ATG9* (AT2G31260) in WT and *dapat* plants exposed to prolonged darkness (Figure 4A-C). In *dapat*, *ATG7* and *ATG8b* transcripts were rapidly and strongly induced by day 3, reaching approximately 8-fold and 28-fold increases, respectively, and exceeding the response observed in WT. By day 7, however, transcript abundance declined in *dapat*, converging with WT levels (Figure 4A, B). In striking contrast, *ATG9* remained consistently repressed in *dapat* throughout the treatment, while WT displayed a progressive increase that peaked at day 7; notably, reduced *ATG9* expression was already evident in the mutant before darkness imposition (Figure 4C). Together, these patterns indicate that lysine biosynthesis impairment in *dapat* triggers an early and selective activation of autophagy-related pathways, whereas *ATG9* induction remains constrained.

**Figure 4.**
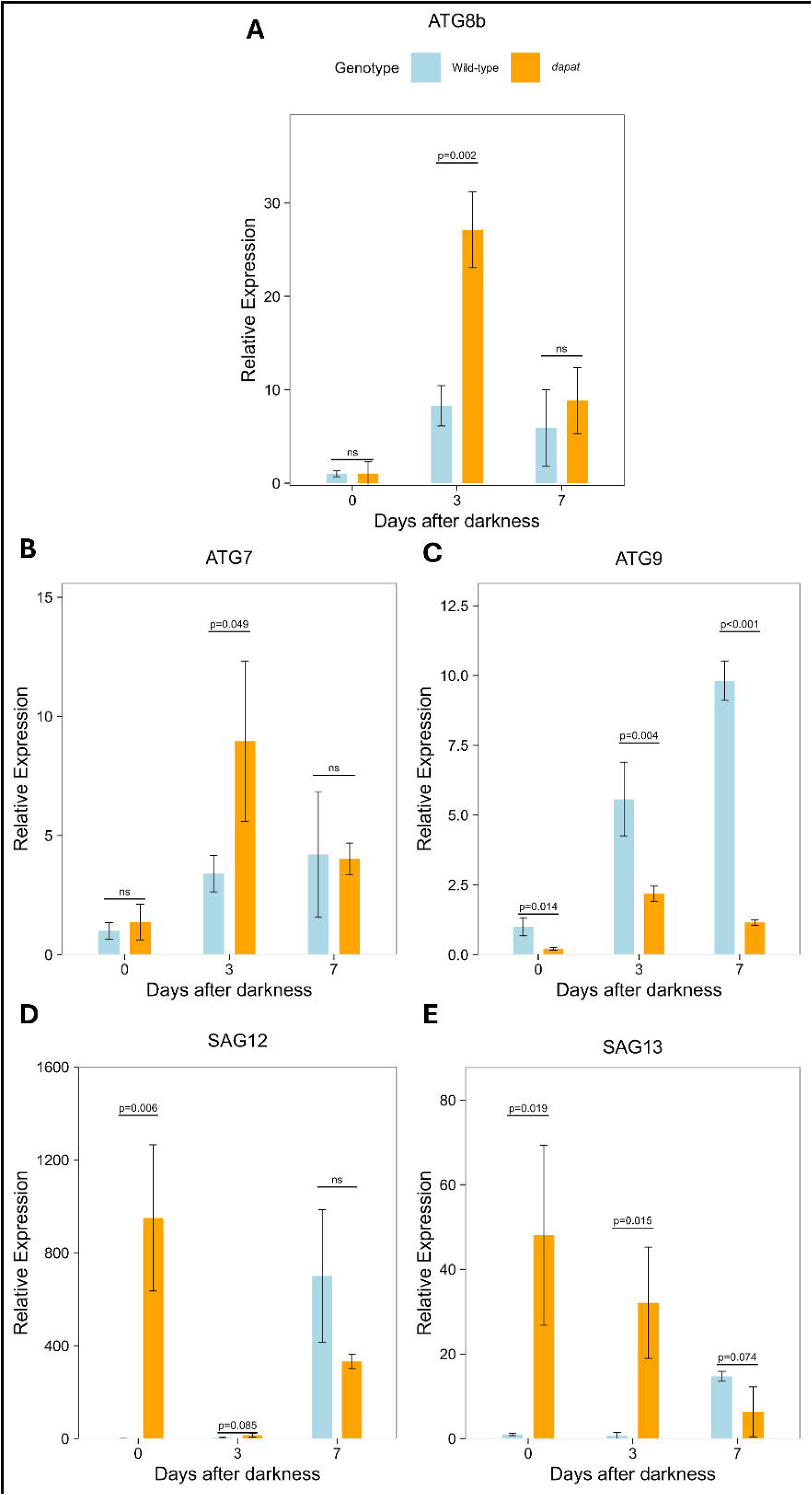
DAPAT mutation triggers basal leaf senescence and intensifies selective autophagic activity during extended darkness. Transcript levels of (A) *ATG8b*, (B) *ATG7*, (C) *ATG9,* (D) *SAG12*, and (E) *SAG13* in five-week-old *Arabidopsis thaliana* wild-type and *dapat* plants grown under short-day (8h/16h light-dark) conditions, 150 μmol photons m ²s ¹, 20°C day/18°C night and subjected to extended darkness. Data represent transcript abundance expressed as mean ± SEM of three biological replicates. P values indicate significant differences compared to wild-type at each time point by two-tailed Student’s t-test. Expression levels were normalized using the reference genes *UBC9* and *UBC21*. The reference group for the ΔΔCt method used was WT: day 0.

### Lysine biosynthesis deficiency locks plants into a constitutive starvation-like state

To determine whether the accelerated senescence and impaired recovery of mutant plants under continuous darkness depend on the preceding photoperiod, we grew WT and *dapat* plants for four weeks under neutral-day (12h/12h light–dark; ND) before subjecting them to darkness imposition (Figure 5A), followed by a four-day recovery period (Figure 5D). Strikingly, even before stress, ND-grown mutants already displayed lower *Fv/Fm* values, and this difference increased during the treatment, with *dapat* reaching nearly three-fold lower values by day 7 (Figure 5B), confirming increased PSII vulnerability independent of photoperiod. Remarkably, both genotypes recovered after transfer back to optimal growth conditions under a 12 h/12 h light–dark photoperiod (Figure 5D). Dark respiration rates in WT and *dapat* leaves remained largely similar throughout treatment. No significant differences were detected between genotypes at day 0, and only a modest decline was observed by day 3, with rates returning to comparable values by day 7 (Figure 5C).

**Figure 5.**
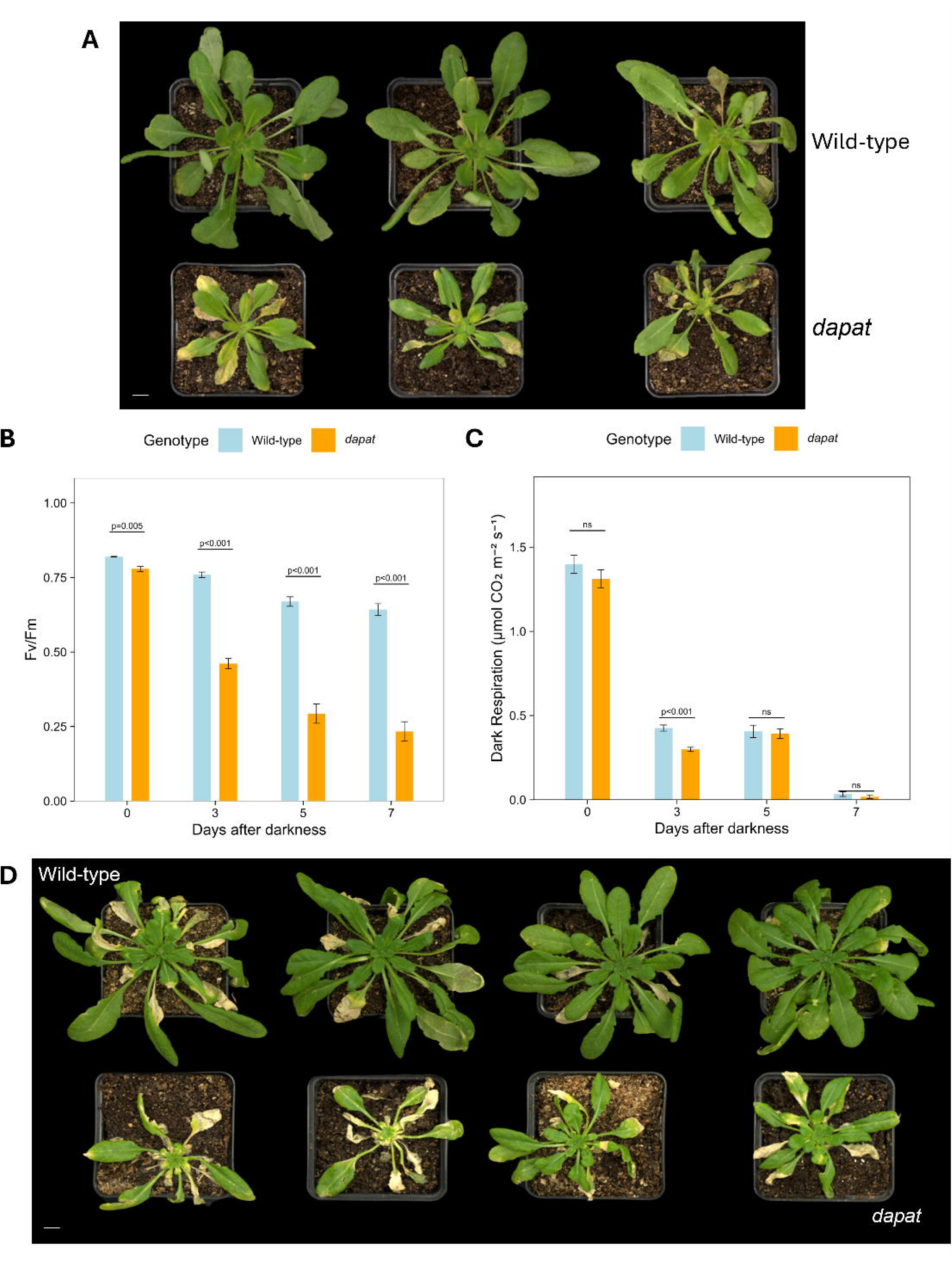
Lysine biosynthesis disruption impairs photochemical and respiratory responses to extended darkness. (A) Phenotypes of five-week-old *Arabidopsis thaliana* wild-type (WT) and *dapat* mutant plants grow under neutral-day (12h/12h light-dark) conditions, (20°C day/18°C night) with a light intensity of 150 µmol m ² s ¹, followed by nine days of extended darkness. (B) Maximum quantum efficiency of photosystem II (*Fv/Fm*) in leaves of WT and *dapat* plants during seven days of darkness. Data represents the mean ± SEM of eight biological replicates (two leaves per plant). (C) Dark respiration rates in leaves of plants exposed to seven days of darkness. Values are presented as the mean ± SEM of five independent biological replicates. P values denote significant differences between genotypes at each time point by two-tailed Student’s t-test. (D) Phenotypes of *Arabidopsis thaliana* wild-type (WT) and *dapat* mutant plants grow under neutral-days (12h/12h light-dark) conditions. Plants were subjected to nine days of continuous darkness followed by a 4-day recovery period under the initial growth conditions. White scale bars at the left bottom of Fig. 5A and Fig. 5D represent 1 cm.

Following the recovery of survival capacity in mutant plants after 9 days of induced darkness, we asked whether impaired lysine biosynthesis prevents proper metabolic resetting after carbon starvation, leading to persistent activation of alternative respiratory pathways and incomplete restoration of carbohydrate homeostasis after recovery. In the short-day experiment described above, however, the severe senescence already evident at the end of the treatment greatly limited the harvest of plant material, precluding a more extensive metabolic analysis. To further address this, we analyzed the metabolic impact of the treatment under ND photoperiod.

Organic acid dynamics showed a differential accumulation of TCA cycle intermediates, including 2-oxoglutarate (2-OG), citrate, isocitrate, malate, and succinate in both genotypes. This pattern was markedly pronounced in *dapat* on day 5, reflecting altered respiratory demand during stress. Notably, this metabolic behavior persisted after recovery, consistent with the phenotype already evident from day 0 (Figure 6). In contrast, sugar depletion showed the opposite trend, with the darkness treatment promoting accelerated consumption of sucrose and soluble sugars, including glucose and fructose. Raffinose levels also declined markedly in response to stress, particularly in *dapat*, suggesting an early disruption of raffinose family oligosaccharide (RFO) metabolism that persisted throughout both stress and recovery (Figure 6). After the 4-day recovery period, WT plants showed a greater capacity to restore metabolic homeostasis, with all analyzed metabolites returning to control levels (Figure 6). In *dapat*, most metabolites also returned to baseline, except for proline, glutamate, glutamine, and glycine, which remained elevated. To distinguish constitutive metabolic differences in the mutant from time-dependent responses to darkness, metabolite abundances in *dapat* were normalized to the corresponding WT levels at each time point (Figure S4). This analysis revealed a consistent pattern of organic and amino acids accumulation across all conditions, indicating a persistent metabolic state that is largely independent of stress duration.

**Figure 6.**
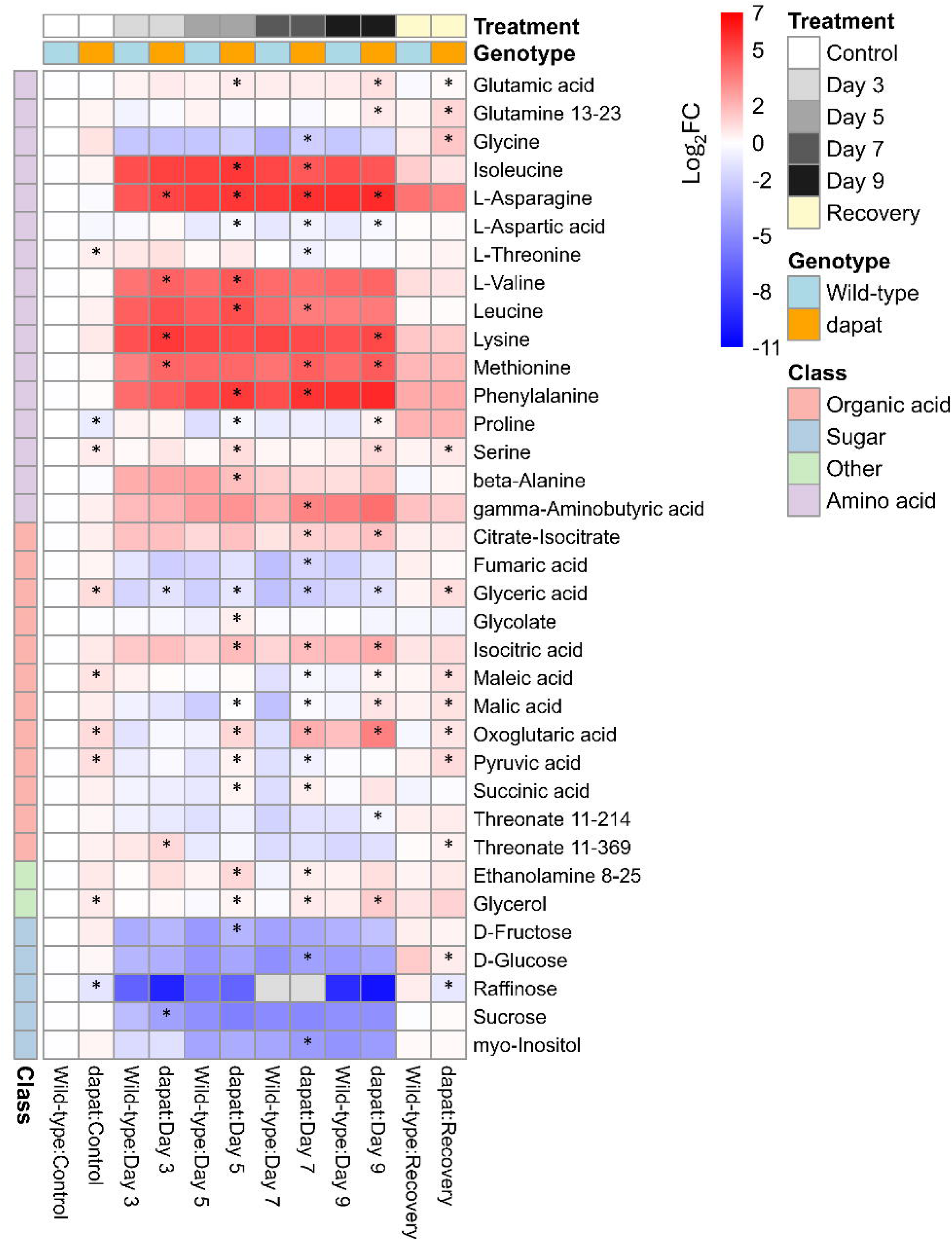
DAPAT mutation elicits global metabolic reprogramming during dark-induced senescence. Five-week-old *Arabidopsis thaliana* wild-type (WT) and *dapat* mutant plants grown under neutral day (12h/12h light-dark) conditions were exposed to continuous darkness for 3, 5, 7, or 9 days, followed by a recovery period under light. Each cell represents the log fold change in metabolite levels normalized to WT:Control (day 0). Colors represent the log fold change: red indicates enrichment, blue indicates depletion, and white represents no change. Genotype colors are light blue (WT) and orange (*dapat*). Treatments colors are represented as white (control), light gray (day 3), medium gray (day 5), dark gray (day 7), black (day 9) and yellow (recovery period). Metabolite classes are indicated in the right bar: green for amino acids, pale yellow for organic acids, red for sugars, and light purple for other metabolites. Values represent the mean ± standard error of five independent samples. Statistical significance versus WT shown by asterisks; each group normalized to its corresponding WT control day 0 (two-tailed Student’s t-test; P < 0.05).

To further reinforce the constitutive metabolic priming phenotype triggered by the DAPAT mutation under energy limitation, we performed principal component analysis (PCA) to integrate the metabolomic datasets (Figure S5). PC1 (53.4% of the variance) mainly reflected carbohydrate and organic acid depletion–restoration dynamics, with late-darkness samples shifting to negative scores and recovery samples returning to positive values. PC2 (18.9% of the variance), driven by amino acids and polyols (including glycerol, serine, succinate, 2-OG), clearly separated *dapat* from WT plants under stress conditions. WT samples showed a clear reduction in sugars without a strong increase in TCA cycle intermediates, whereas *dapat* showed reduced carbohydrates together with higher organic acids and amino acids levels, suggesting a genotype-specific metabolic rerouting.

During recovery, both genotypes shifted toward positive PC1 values, pointing to carbohydrate restoration. However, *dapat* mutant retained higher PC2 scores, consistent with sustained TCA cycle activation and continued amino acid oxidation. Notably, WT under recovery clustered closely with unstressed *dapat*, indicating that stress-induced metabolic reprogramming in WT plants converges toward a constitutive metabolic configuration already present in *dapat* under control conditions. In summary, the integrative metabolomic analysis indicates that the DAPAT mutation imposes a constitutive stress-like metabolic state that closely resembles the transient starvation response adopted by WT plants during energy privation. These findings extend previous reports by showing that lysine biosynthesis deficiency not only primes a stress-like metabolic state, but also impairs metabolic resetting after extended darkness, preventing full restoration of homeostasis.

### Lysine deficiency impairs metabolic reset after carbon starvation

Based on the persistent stress-like metabolic state observed in *dapat* plants, we examined whether altered energy signaling targets were also affected by lysine biosynthesis disruption, we assessed the transcriptional dynamics of the catalytic subunits of the Sucrose non-fermenting-related kinase 1 (SnRK1) complex KIN10 (AT3G01090) and KIN11 (AT1G32290). Under control conditions (day 0), *dapat* plants exhibited constitutively reduced *KIN10* transcript levels (∼2 -fold lower than WT) (Figure 7A), suggesting compromised basal energy sensing. Following the return to light, *KIN11* expression was significantly repressed in *dapat* (∼1.3 -fold reduction) (Figure 7B), contrasting sharply with the WT plants, which restored transcript levels toward baseline. Together, these data support the hypothesis that impaired lysine biosynthesis compromises the transcriptional recovery of SnRK1-related energy signaling, which may underlie the incomplete metabolic restoration observed in *dapat* mutant plants.

**Figure 7.**
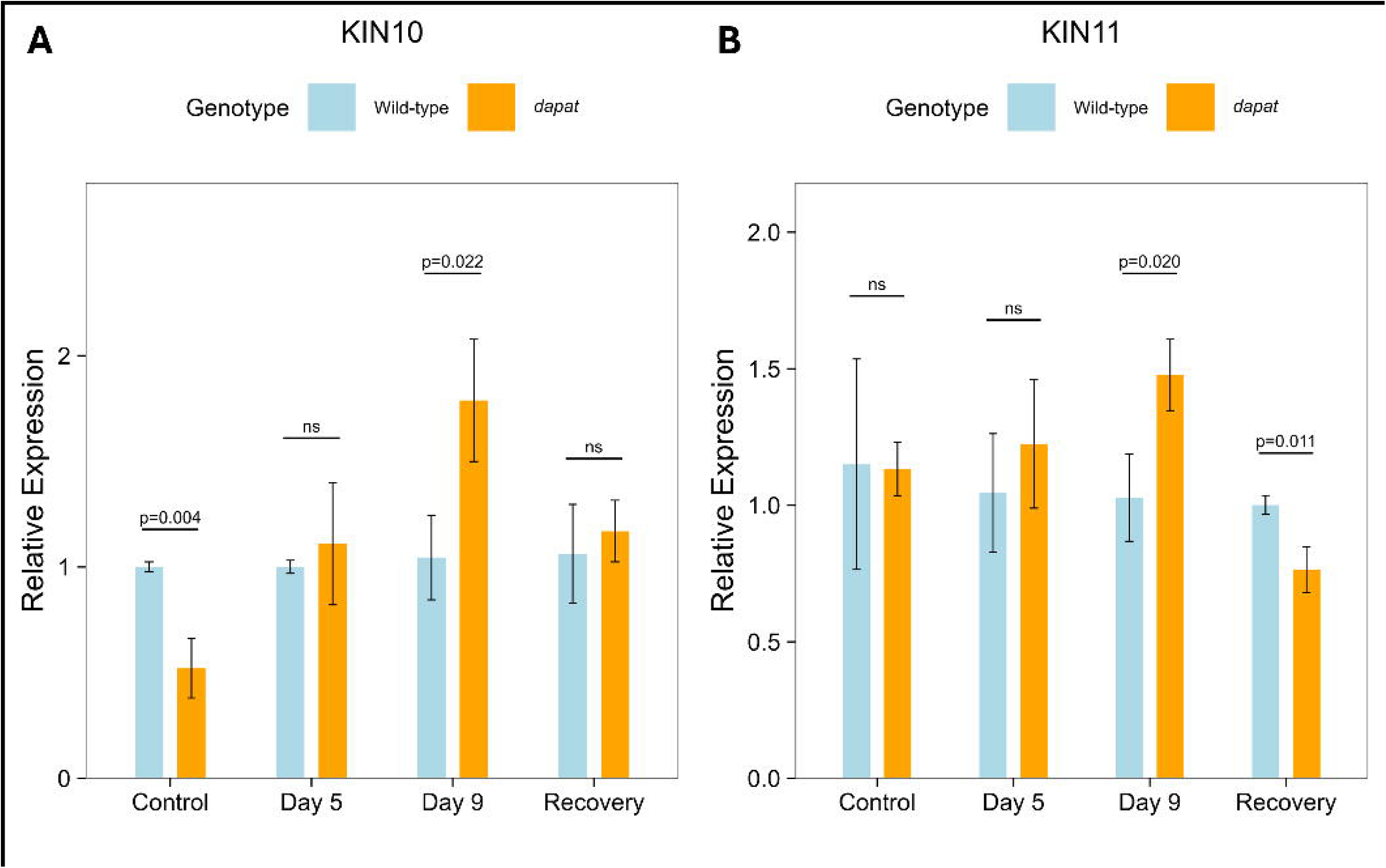
Transcriptional dynamics of *SnRK1* subunit during darkness starvation in *dapat*. Transcript levels of *KIN10* (AT3G01090) and *KIN11* (AT3G29160) in *Arabidopsis thaliana* wild-type (WT) and *dapat* mutant plants grown under neutral-day (12h/12h light-dark) conditions, 150 μmol photons m ²s ¹, 20°C day/18°C night, and subjected to nine days of darkness followed by four days of recovery. Data represent transcript abundance expressed as mean ± SEM of three biological replicates. P values indicate significant differences versus wild-type at each time point (two-tailed Student’s t-test. Expression levels were normalized using the reference genes *MON1* and *TIP41*. The reference group for the ΔΔCt method used was WT: day 0.

**Figure 8.**
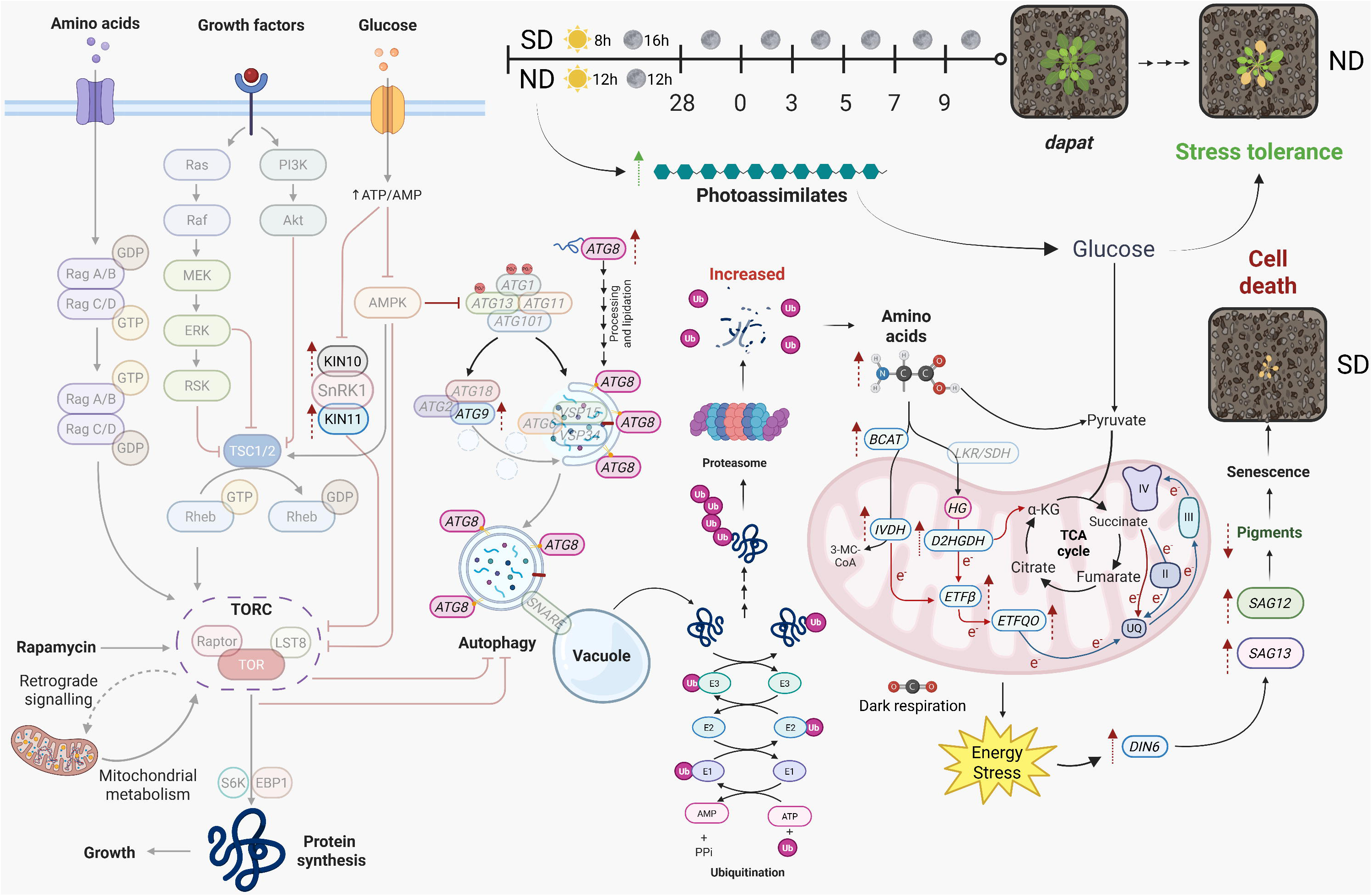
Metabolic and molecular model showing the involvement of lysine biosynthesis in the responses of *Arabidopsis* to extended darkness. Four-week-old *Arabidopsis thaliana* wild-type (WT) and *dapat* mutant plants grown under short-day (8h/16h light-dark, SD) and neutral-day (12h/12h light-dark, ND) conditions were subjected to extended darkness for 9 days, with samples collected at days 0, 3, 7, and 9, as well as during the subsequent recovery period. The model illustrates interconnected pathways regulating stress responses during darkness. The left panel depicts signaling cascades initiated by amino acids, growth factors, and glucose, which activate the Target of rapamycin complex (TORC) pathway comprising TOR (target of rapamycin), Raptor (regulatory-associated protein of TOR), LST8 (lethal with sec thirteen 8) and the SNF1-related kinases KIN10 and KIN11. Central pathways show autophagy regulation through ATG (autophagy-related) proteins: ATG1, ATG3, ATG7, ATG8, ATG9, ATG10, ATG11, ATG13, ATG101, and their interactions with AMPK (AMP-activated protein kinase). The right panel displays metabolic alterations, including upregulation (↑) of branched-chain amino acid transaminase (*BCAT-2*), isovaleryl-CoA dehydrogenase (*IVDH*), D-2-hydroxyglutarate dehydrogenase (*D2HGDH*), electron transfer flavoprotein beta subunit (*ETF*β), ETF: ubiquinone oxidoreductase (*ETFQO*), and the senescence-associated genes *SAG12* and *SAG13.* Non-transparent gene symbols indicate direct evaluation in this study, while transparent symbols represent additional components of the depicted pathways. Extended darkness induces energy stress, alters amino acid metabolism, enhances autophagy, and ultimately leads to senescence or stress tolerance depending on genotype, with *dapat* mutant plants showing greater stress tolerance under ND conditions.

## 4. Discussion

Lysine metabolism homeostasis has increasingly been recognized as an important determinant of plant stress resilience (Yang *et al*., 2020, Arruda & Azevedo, 2020), particularly through its contribution in alternative respiratory pathways during carbon limitation (Araújo *et al*., 2011; Hildebrandt, 2018). Consistent with this interpretation, the hypomorphic *dapat* line, which retains only ∼10% of DAPAT activity, displays a chronic energy-deprivation phenotype even under control conditions, marked by severe defects in germination, growth, photosynthesis, major alterations in carbon-nitrogen (C/N) balance and induction of stress-responsive genes (Cavalcanti *et al*., 2018; Gouveia *et al*., 2026). Although *dapat* was originally identified as a constitutive defense mutant formerly (named *agd2-1*), characterized by enhanced resistance to *Pseudomonas syringae*, elevated salicylic acid levels, and developmental abnormalities (Rate & Greenberg, 2001), our findings extend this view by demonstrating that DAPAT fulfils central metabolic roles that go beyond immunity, pointing lysine biosynthesis as contributors to basal metabolic homeostasis rather than functioning exclusively under stress conditions (Cavalcanti *et al*., 2018; Neves *et al*., 2024; Gouveia *et al*., 2026), further supported by the embryonic lethality of knockout mutants (Rate & Greenberg, 2001; Song *et al*., 2004).

Here, we show that impaired lysine biosynthesis due to reduced DAPAT activity does more than limit substrates for respiration. Instead, it creates a constant energy-limited metabolic state that reduces metabolic flexibility during extended darkness. To test this, WT and *dapat* plants grown under short-day (SD) conditions were subjected to nine days of continuous darkness, following the experimental design demonstrated by Barros *et al*. (2017). Severe carbon deprivation revealed a pronounced survival defect in the mutant (Figure 1A, B and Figure 5). Even prior to dark treatment, *dapat* plants already displayed reduced chlorophyll content and lower *Fv/Fm* (Figure 1C and Figure S3), indicating compromised photosystem II efficiency. Chronic chlorophyll depletion likely accelerates carbon–nitrogen remobilization from senescing leaves through degradation of major nitrogen pools, including chlorophyll-binding proteins and RuBisCO, thereby releasing phototoxic intermediates that must be neutralized (Zhu *et al*., 2022; Otegui *et al*., 2005). In line with this early decline in photosynthetic capacity, autophagy related and senescence associated transcriptional targets showed basal induction in *dapat* plants, in parallel with enhanced protein turnover and amino acid accumulation even before stress (Figure 2). This pattern supports the previous reported disturbance in C/N homeostasis (Cavalcanti *et al*., 2018; Neves *et al*., 2024; Gouveia *et al*., 2026).

Darkness exposure further revealed genotype-specific differences in the expression of senescence-related genes. In *dapat*, *SAG12* and *SAG13* were induced early but transiently, whereas WT showed a delayed peak in SAG expression at day 7 (Figure 4D-E), suggesting an out-of-phase senescence response. However, this premature activation did not translate into improved metabolic resilience. This interpretation is reinforced by the differential expression of autophagy-related genes in the mutant. Although *ATG7* and *ATG8b* were basal activated in *dapat* (day 3) and reached WT levels by day 7 (Figure 4A-B), *ATG9* remained repressed (Figure 4C). The premature activation of *ATG7* and *ATG8b* is consistent with autophagic priming (Li & Vierstra, 2012; Avin-Wittenberg *et al*., 2015, 2019; Wang *et al*., 2018; Zhu *et al*., 2022), whereas sustained repression of *ATG9* suggests impaired autophagosome maturation, likely limiting autophagic flux and exacerbating energy imbalance in DAPAT-deficient plants (Zhuang *et al*., 2017). Previous transcriptome analyses by microarray had already reported a basal induction of ATG genes in *dapat* mutant, likely triggered by DAPAT disruption (Cavalcanti *et al*., 2018).

This phenotype of catabolic pathway activation becomes particularly evident at the transcriptional level, especially in genes operating downstream of lysine and BCAA catabolism. In *dapat*, *BCAT-2* was already induced at day 0 but showed only limited further activation during extended darkness (Figure 3A), suggesting that impaired lysine biosynthesis constraints sustained BCAAs catabolism and, consequently, the energy supply required under SD conditions. Despite this activation, the muted response points to a pre activated metabolic state unable to properly respond to energy deprivation. By contrast, *IVDH* transcript levels, which channels BCAAs through an alternative oxidative route, remained strongly induced in *dapat* throughout darkness (Figure 3B), reinforcing a compensatory activation of alternative oxidative routes to support respiration when lysine synthesis is compromised (Araújo *et al*., 2010, 2011). In WT, *IVDH* induction was attenuated under carbon privation, reflecting a more balanced metabolic adjustment. Additionally, *D2HGDH* was transiently upregulated in WT during dark stress but only at day 0 in *dapat* plants (Figure 3C), supporting the premature yet unstained metabolic priming. Examination of *ETF*β and *ETFQO* transcripts under stress further indicate genotype-specific differences in alternative respiration dynamics (Figure 3D–E). While *ETF*β was induced in WT but repressed in *dapat*, *ETFQO* was strongly upregulated in the mutant, suggesting compensatory activation of the oxidoreductase to maintain electron flux despite impaired upstream substrate supply (Ishizaki *et al*., 2005, 2006; Peng *et al*., 2015). Extending previous models of amino acid-mediated stress adaptation, our findings reveal that DAPAT deficiency does not simply alter lysine catabolism as an alternative respiratory route. In this context, *dapat* plants appear to remain in a low-plasticity state that limits the re-establishment of carbon balance after stress.

### Photoperiod shapes dark survival in *dapat* mutant plants

The transcriptional analysis of SnRK1-related components suggested altered energy sensing, with *KIN10/KIN11* expression pointing to a starvation-responsive transcriptional state that coordinates catabolic activation and remobilization. Consistent with the observed raffinose depletion (Figure 6), *dapat* mutant plants exhibited constitutive repression of *KIN10* (Figure 7A, day 0), suggesting compromised basal energy sensing and a potentially pre-existing metabolic imbalance (Cavalcanti *et al*., 2018; Neves *et al*., 2024). Under normal conditions, *KIN10* expression increases at nightfall to promote carbon remobilization and repress anabolic processes (Baena-González *et al*., 2007; Wurzinger *et al*., 2016; Pedrotti *et al*., 2018; Peixoto & Baena-González, 2022). During stress, dapat plants displayed a compensatory upregulation of both *KIN10* and *KIN11*, peaking at day 9 (Figure 7). In addition, although *KIN10* expression returned to WT levels upon re-illumination (Figure 7A), whereas *dapat* plants failed to fully re-establish metabolic homeostasis (Figure 6 and Figure S5). Together, this photoperiod-dependent recovery profile highlights a sustained metabolic reprogramming imposed by the DAPAT mutation and reveals incomplete restoration of energy balance following carbon limitation. Importantly, our findings extend beyond the established role of lysine as a respiratory substrate (Araújo *et al*., 2010, 2011; Barros *et al*., 2017; Cavalcanti *et al*., 2018). Rather than simply fueling mitochondrial respiratory metabolism, a well-established principle, our findings unveil a more fundamental regulatory function for the pathway, demonstrating that lysine *de novo* biosynthesis and their homeostasis exerts a broader regulatory role in maintaining metabolic flexibility during energy stress, consistent with the emerging roles of amino acids as both metabolic substrates and signaling molecules in plants (Heinemann & Hildebrandt, 2021; Bady *et al*., 2026).

## 5. Conclusions

This work supports a broader role for lysine homeostasis tightly integrated with energy and metabolic sensing that underlies plant survival. Beyond supplying proteinogenic lysine, DAPAT-mediated biosynthesis emerges as a central node coordinating nitrogen remobilization, mitochondrial substrate feeding, and starvation responses. Strikingly, this metabolic vulnerability is photoperiod-dependent: short-day growth imposes pre-existing carbon constraints that, when combined with prolonged darkness, push *dapat* mutant plants beyond a survival threshold. In contrast, neutral-day conditions permit sufficient photoassimilate accumulation to sustain survival despite reduced DAPAT activity, demonstrating that lysine biosynthesis deficiency creates a state of metabolic fragility shaped by the growth photoperiod.

These findings challenge the traditional view that amino acid catabolism during starvation primarily reflects substrate availability. Instead, they indicate that amino acid metabolism is tightly coupled to energy-sensing networks, and that lysine biosynthesis itself contributes to regulating these responses. When disrupted, this coordination fails, cascading through autophagy, respiration, and senescence programs. Lysine biosynthesis thus emerges as a key determinant of metabolic plasticity, shaping the plant’s capacity to enter, sustain, and exit starvation-induced metabolic states. By positioning DAPAT as a regulatory node at the interface of carbon and nitrogen metabolism, this work expands our mechanistic understanding of how primary metabolism integrates with stress adaptation and d highlights lysine-dependent regulatory networks as potential targets for improving crop resilience under resource limitation.

## Acknowledgments

This research was funded by the National Council for Scientific and Technological Development (CNPq Brazil, Grant 407276/2021 1 and 151020/2024-8), the National Institute of Science and Technology in Plant Stress Physiology (INCT Plant Stress Physiology/CNPq Brazil, Grant 406455/2022 8), and the FAPEMIG (Foundation for Research Assistance of the Minas Gerais State, Brazil, Grant RED 00060 23). Scholarships to DGG from CAPES PRINT and research fellowships from CNPq Brazil to ANN and WLA are also acknowledged. APMW acknowledges funding by the Deutsche Forschungsgemeinschaft Cluster of Excellence for Plant Sciences (CEPLAS) under Germany’s Excellence Strategy EXC-2048/1 under project ID 390686111.

## 6. Author Contributions

D.G.G. and W.L.A. designed the research; D.G.G performed the research with the support of A.N-N., A.P.M.W., and W.L.A.; W.E.B.B., P.W., A.O.M., S.F., B.P.Z., J.H.F.C., and A.P.M.W. contributed new reagents/analytic tools; D.G.G., A.N-N., A.P.M.W., and W.L.A. analyzed the data; and D.G.G., A.R.F., A.N-N., A.P.M.W., and W.L.A. wrote the article with input from all the others.

## 8. Data availability

The data that support the findings of this study are available from the corresponding author upon reasonable request.

## Supplementary Tables Legends

**Table S1.** Primers used in the RT-qPCR analyses performed in this study.

## Supplementary Figures Legends

**Figure S1. Genetic complementation of the *dapat* mutant restores the wild-type phenotype.** *Arabidopsis thaliana* wild-type (WT), *dapat* mutant, and complemented (DAPAT gene F3 generation) plants were grown under neutral-day (12h/12h light-dark) conditions, at 150 μmol photons m ² s ¹. Two independent complementation lines are shown: plant 1 and plant 2 for the pUTKAN HA-tagged vector, and plant 1 for the pUTKAN vector.

**Figure S2. The *dapat* mutant is unable to recover from extended darkness in short-day-grown plants.** Plants were initially grown under short-day (8h/16h light-dark) conditions, 150 μmol photons m ²s ¹, 20°C day/18°C night for five weeks before exposure to continuous darkness. (A) Phenotype of five-week-old *Arabidopsis thaliana* wild-type and *dapat* mutant plants after 9-days of dark treatment (DAD), and (B) phenotype following a subsequent 7-day recovery period.

**Figure S3. Extended darkness accelerates chlorophyll depletion in leaves of the *dapat* mutant.** (A) Chlorophyll a, (B) chlorophyll b and (C) chl a/b ratio in leaves of five-week-old *Arabidopsis thaliana* wild-type and *dapat* mutant plants during extended darkness. Plants were grown under short-day (8h/16h light–dark) conditions 150 μmol photons m ²s ¹, 20°C day/18°C night. Asterisks indicate significant differences versus wild type at each time point (Tukey’s HSD test; P < 0.05). Data represents the mean ± standard of five biological replicates.

**Figure S4. Metabolic remodeling under extended darkness is intensified by the *dapat* mutation.** Five-week-old *Arabidopsis thaliana dapat* mutant plants grown under neutral-day (12h/12h light-dark) conditions were exposed to continuous darkness for 3, 5, 7, or 9 days, followed by a recovery period under light. Each cell represents the log fold change in metabolite levels normalized to wild-type: red indicates enrichment, blue indicates depletion, and white represents no change. Treatment colors are represented as white (control), light gray (day 3), medium gray (day 5), dark gray (day 7), black (day 9) and yellow (recovery period). Metabolite classes are indicated in the right bar: green for amino acids, pale yellow for organic acids, red for sugars, and light purple for other metabolites. Values represent the mean ± standard error of five independent samples. Statistical significance versus WT shown by asterisks; each group normalized to its corresponding WT control day 0 (two-tailed Student’s t-test; P < 0.05).

**Figure S5. Metabolite profiling by PCA highlights distinct responses of WT and *dapat* to extended darkness and recovery.** Principal component analysis (PCA) score plot of metabolite profiles from *Arabidopsis thaliana* wild-type (WT) and *dapat* mutant plants under control, extended darkness and recovery treatments. PC1 and PC2 explain 53.4% and 18.9% of the total variance, respectively. Overlaid loading vectors highlight top 5 most strongly metabolites contributing to PC1 (red arrows: myo-inositol, sucrose, *D*-fructose, fumaric acid, *L*-valine) and PC2 (blue arrows: glycerol, succinic acid, 2-oxoglutarate, serine). The distinct separation along PC1 reflects shifts in carbohydrate and organic acid metabolism under extended darkness, whereas PC2 captures variation in amino acids and polyol profiles associated with genotype and stress response.

