## Supplementary figures and images for "Impaired lysine biosynthesis primes constitutive energy stress and dark-stress sensitivity"

### Figure S1

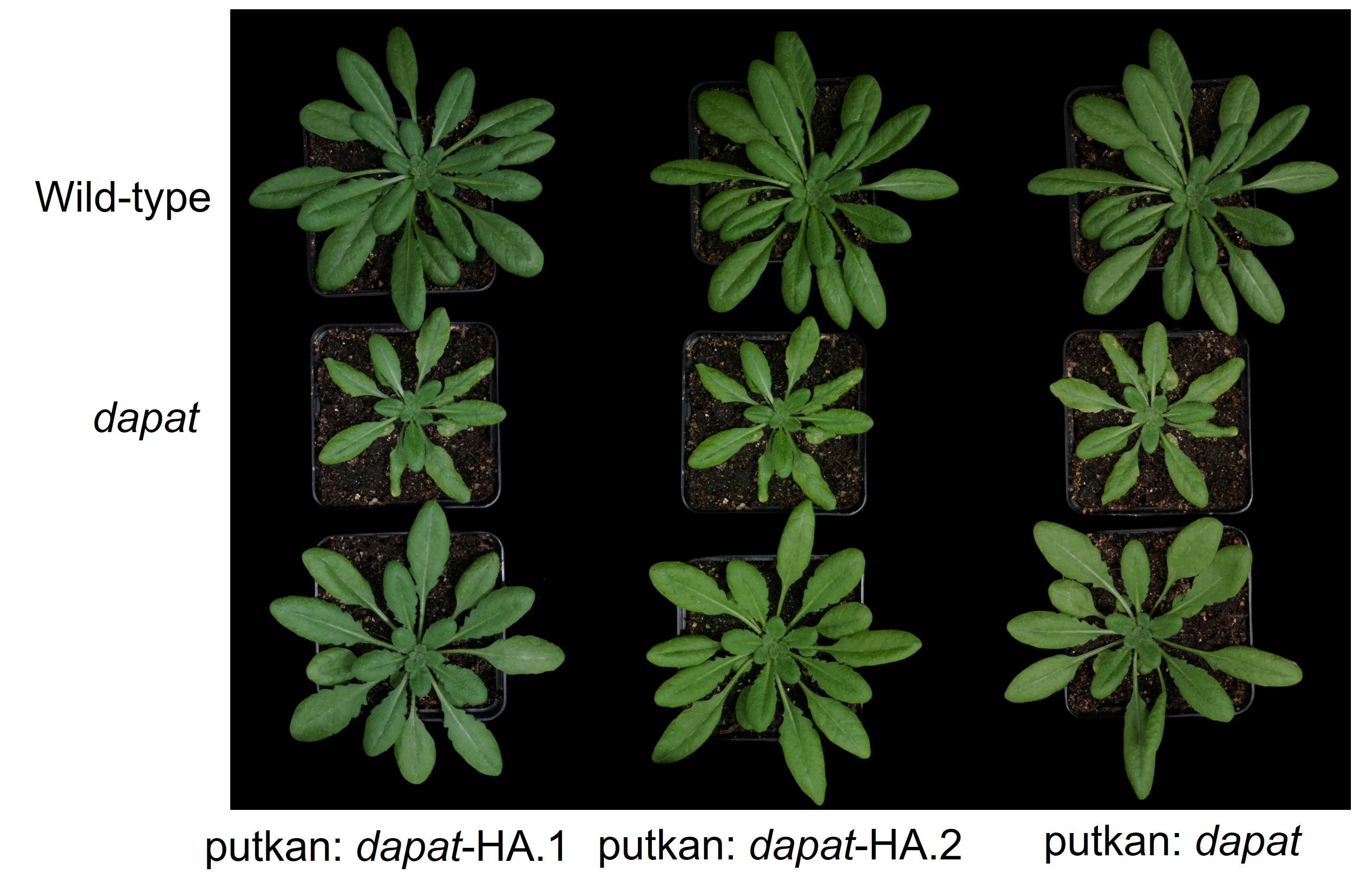

### Figure S2

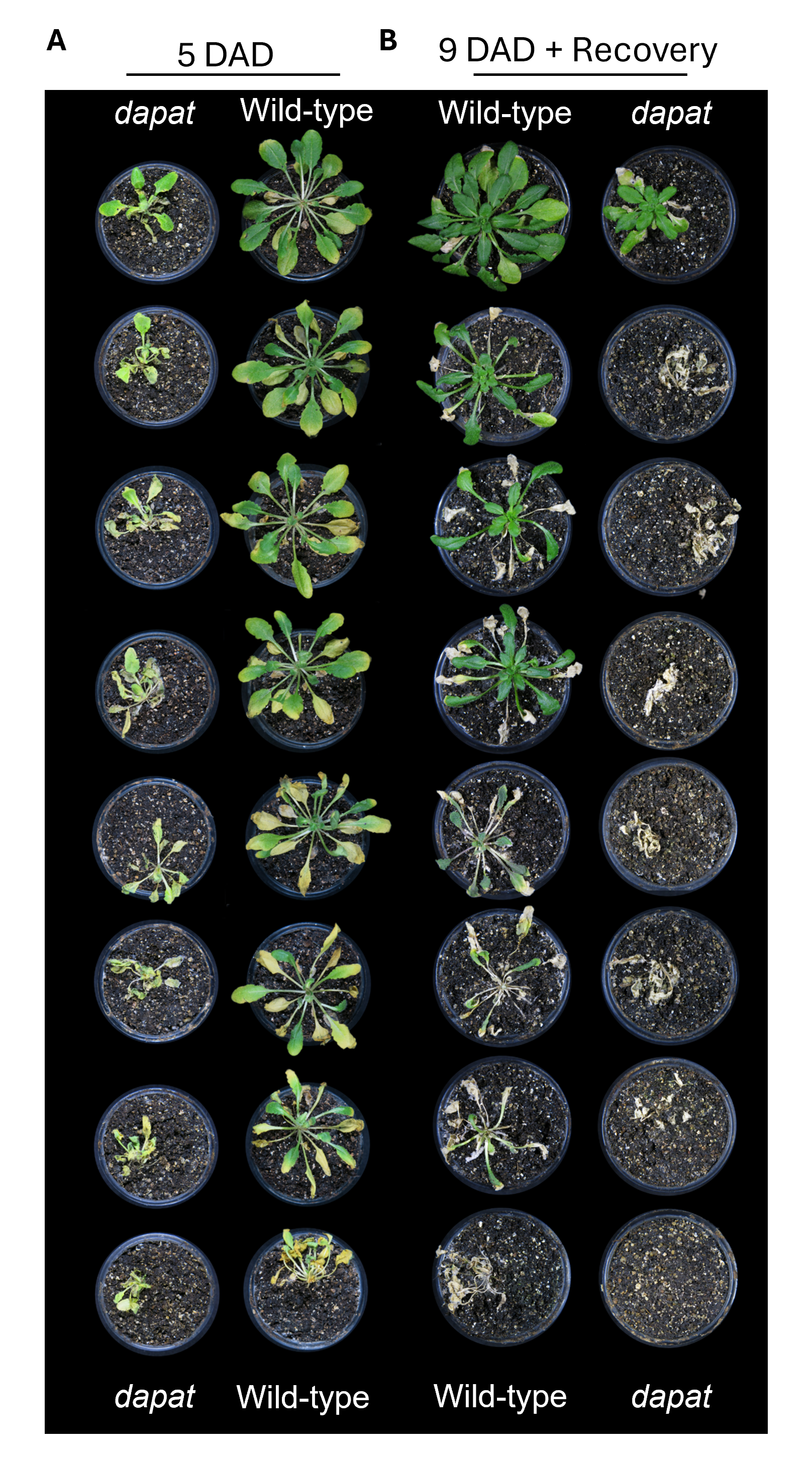

### Figure S3

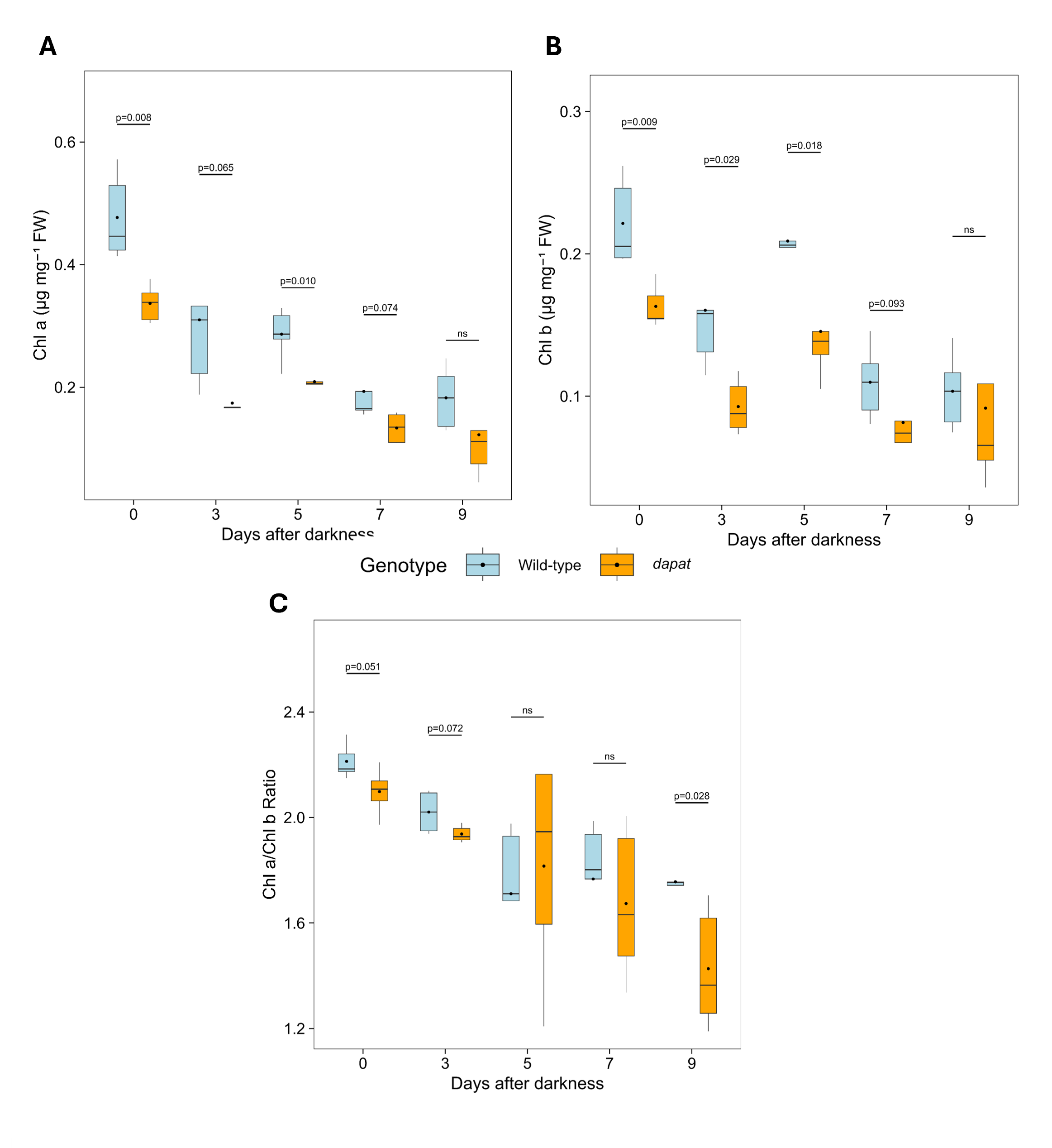

### Figure S4

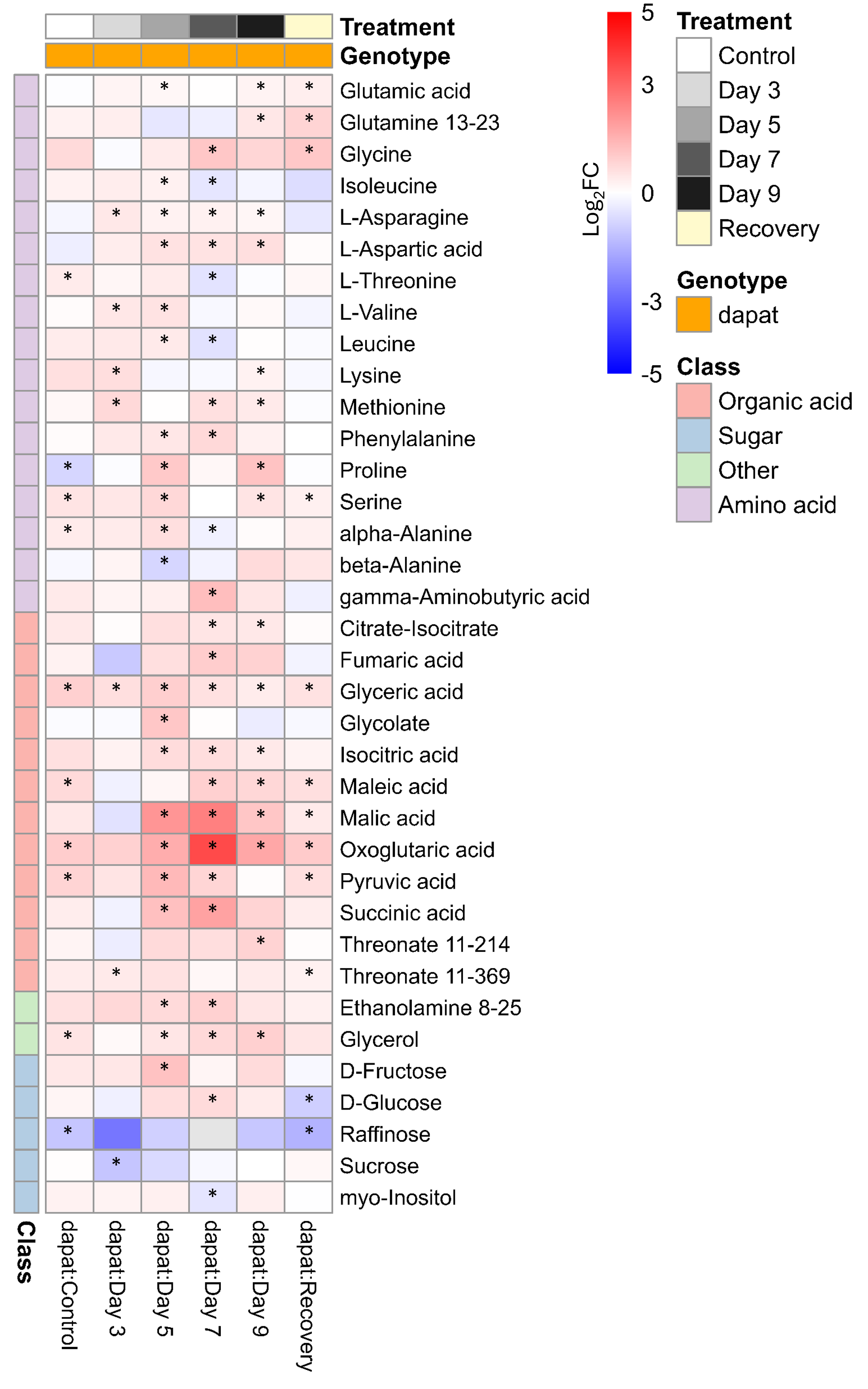

### Figure S5

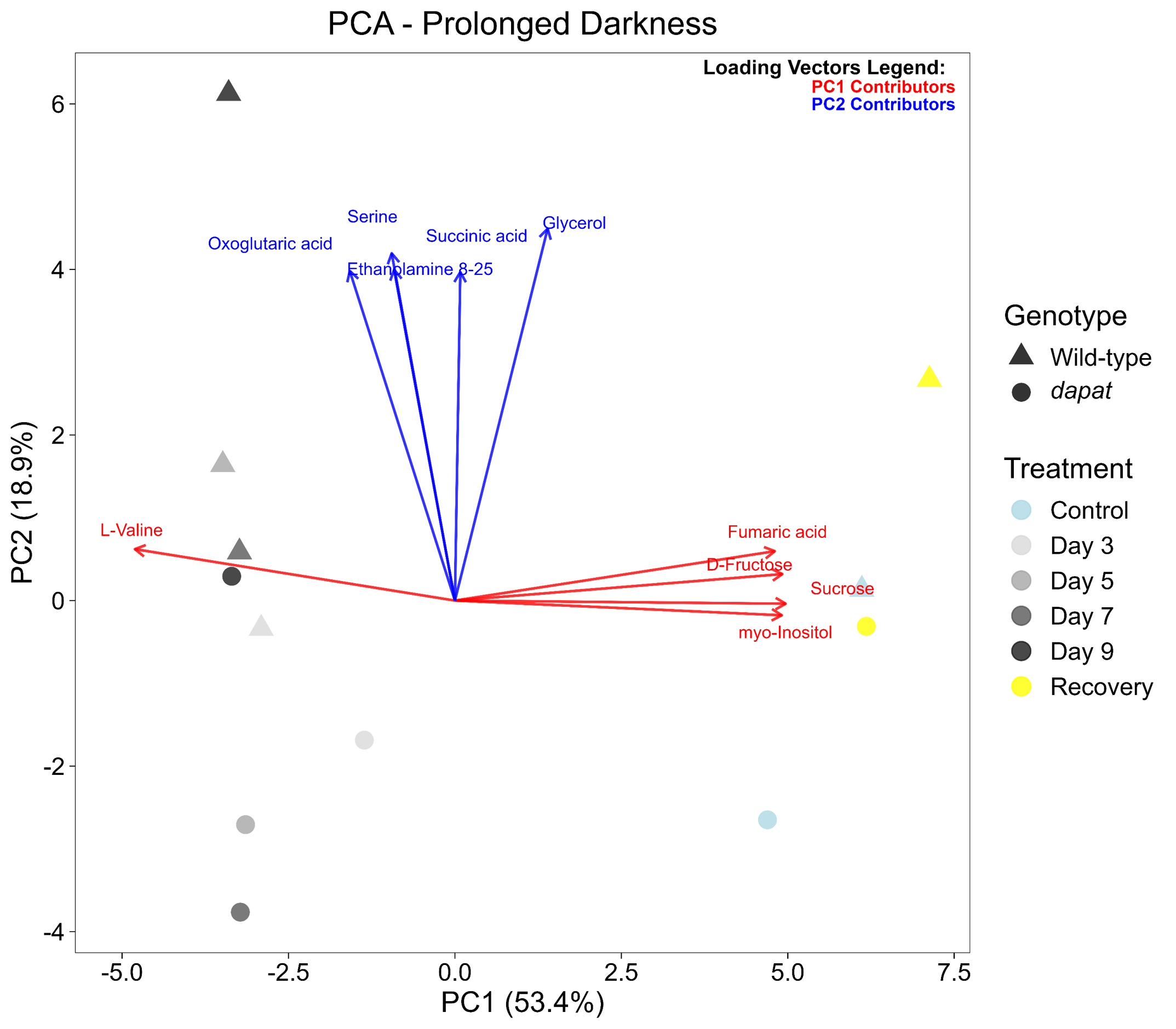
