## Supplementary material for "Impaired lysine biosynthesis primes constitutive energy stress and dark-stress sensitivity": Table S1

**Supplementary Table S1** - Primers used for RT-qPCR assessment of extended darkness gene expression response in *Arabidopsis thaliana* wild-type (WT) and *dapat* mutant plants. *UBC9*, *UBC21, MON1* and *TIP41* were used as reference genes for gene expression normalization. Gene IDs: *DAPAT* (AT4G33680), *BCAT-2* (AT1G10070), *D2HGDH* (AT4G36400), *IVDH* (AT1G04710), *ETFB* (AT2G32360), *ETFQO* (AT2G43400), *SAG12* (AT1G02920), *SAG13* (AT2G39050), *ATG7* (AT5G46830), *ATG8b* (AT4G04620), *ATG9* (AT2G31260), *UBC9* (AT4G27310), *UBC21* (AT5G05250), *MON1* (AT1G49200), *TIP41* (AT4G34270), *KIN10* (AT3G01090), *KIN11* (AT1G32290).

| Gene | Forward Primer | Reverse Primer |
| --- | --- | --- |
| ***DAPAT*** | CTGGAGAAGACTCATGTG | AGATGTTCTCTCTGTGACC |
| ***BCAT-2*** | TCTTTTGTTTCGTCCGGATCA | CAACCGAAGGAGAAGGCATGA |
| ***D2HGDH*** | GAAGCTGTCATATCGGTGGA | TCGTACCCAGTATTGTCTTTGC |
| ***IVDH*** | AATGGGAAAGTTGACCCAAAGGAC | TAAAGCGACCTGCGTTGCTCTC |
| ***ETF Beta*** | CGTTTGCATCGAAGGTTGTGT | CAGTTGTTATCACAGCGGGT |
| ***ETFQO*** | CGAAACTATCGTCCTGCATTCG | TTGCTTCGTGGTCTGCTTTC |
| ***SAG12*** | ACAAAGGCGAAGACGCTACTTG | ACCGGGACATCCTCATAACCTG |
| ***SAG13*** | GGCTTGGGAGAGAACTCAAGA | GCTTCCCCGATGCCTTTAGAG |
| ***ATG7*** | ACGTGGTTGCACCTCAGGATTC | AGGACTCCGACTAAGAGTTCAACG |
| ***ATG8b*** | AGAGTTCCCGTGATTGTGGAA | TCAGCTGGTACAAGATACTTCTTC |
| ***ATG9*** | TTCCACGCAAACCCAATTCT | TGCAACTACAAATCCAGAGACC |
| ***UBC9*** | AAGCATCTGCCTCGACATCT | ATCGATAGCAGCACCTTGGA |
| ***UBC21*** | ACTGCGACTCAGGGAATCTT | TTTCTTAGGCATAGCGGCGA |
| ***MON1*** | AAAGGATTGGGACCCCACAA | TTATCGCCATCGCCTTGTCT |
| ***TIP41*** | GGAAATTCCGGAGCAAGAGCCG | AGAGTAGCTTCCCCCTTTGG |
| ***KIN10*** | CCGCAACCGAACCCAGAATG | CGGGGAGTACCTTCCATGGTC |
| ***KIN11*** | GCAACAGAACACAAAACGATGCTA | GCCACTTGGAACACGGAACC |
